# COMIC: Explainable Drug Repurposing via Contrastive Masking for Interpretable Connections

**DOI:** 10.1101/2025.03.26.645428

**Authors:** Naafey Aamer, Muhammad Nabeel Asim, Andreas Dengel

## Abstract

Many diseases worldwide remain untreated due to the slow and expensive process of drug development. Repurposing existing FDA-approved drugs offers a faster solution, especially with the assistance of artificial intelligence. Despite advancements in AI-driven drug repurposing, current approaches either have lackluster performance or fail to highlight the intricate pathways through which drugs act on diseases. The clinical utility of AI-driven drug repurposing remains constrained by these limitations, particularly for rare and undertreated diseases where data is scarce. To address the need for a precise and explainable predictor, this paper introduces COMIC (**CO**ntrastive **M**asking with **I**nterpretable **C**onnections), a predictor that employs a multi channel architecture consisting of a feature masking branch, which identifies critical drug-disease interaction patterns by extracting the most informative features, and a path masking branch, which highlights relevant biological pathways through which drugs exert their therapeutic effects. Comprehensive evaluation of the COMIC predictor on the PrimeKG knowledge graph (comprising 17,080 diseases, and 4M+ relationships) with nine distinct disease area splits demonstrated a 9.55% average performance improvement over the current state-of-the-art. The practical applicability of the proposed predictor is evaluated on a set of the most recent 30 FDA-approved repurposed drug disease pairs. The COMIC predictor successfully identified 21 of these pairs with high confidence scores. To facilitate real-time drug repurposing investigations, we have developed a publicly available web-based interface for the COMIC predictor (https://drp.opendfki.de/). This application takes disease names as input and returns a ranked list of potential repurposing candidates, along with predicted mechanistic pathways elucidating the drug-disease interactions.

## 1 Introduction

The drug repurposing paradigm encompasses interactions of drugs and disease from two perspectives: indications and contraindications. **Indications** refer to the FDA approved medical uses of a drug, specifically the diseases the drug is approved to treat [1]. These approvals are based on exhaustive clinical research, including clinical trials, real-world evidence, and comprehensive safety and efficacy assessments. Predicting indications is the most significant step for drug repurposing as it encapsulates the potential of a drug to treat a specific disease, forming the primary objective. On the other hand, **Contraindications** are situations where a drug should not be used because it may cause harm to patients with certain diseases [1]. In drug repurposing, understanding contraindications is a secondary step to ensure that a drug being considered for a new use does not pose risks to individuals with specific health conditions. But at the same time, uncovering new indications remains the key objective.

The motivation for finding new indications is grounded in the understanding that drugs often have cascading effects, meaning they can influence multiple biological pathways beyond their primary targets ([2]). Around 30% of FDA-approved drugs have received approval for additional indications beyond their initial intended use with some accumulating over ten new therapeutic applications ([3]). However, many successful drug repurposing efforts have historically resulted from serendipitous discoveries ([4]). These include cases where clinicians observed unexpected benefits when drugs were used for conditions other than their original purpose, such as the use of Metformin to treat polycystic ovary syndrome and the unexpected drug benefits in off-label settings of Thalidomide for multiple myeloma ([5]) ([6]). In contrast to random discoveries, a systematic strategy could enable the repurposing of existing drugs for new therapeutic uses with fewer side effects ([7]). For instance, systematic analysis of drug-protein interaction networks led to the identification of kinase inhibitors like Baricitinib for COVID-19 treatment by targeting both viral entry and inflammation pathways, while network-based screening revealed the potential of antihypertensive drugs like Prazosin for treating post-traumatic stress disorder (PTSD) through shared molecular pathways ([8])([9]).

One of these strategies is the potential matching of drugs to new indications through the analysis of medical knowledge graphs (KGs) ([10]). By evaluating the actions of drugs on cell signaling pathways, gene expression profiles, and disease phenotypes, potential therapeutic candidates can be identified ([2]). The underlying patterns of these actions are learned by geometric deep-learning models, which are trained on extensive medical KGs ([11]). These optimized models can efficiently align disease signatures with drug candidates by analyzing their biological networks ([11]).

While approaches that utilize knowledge graphs (KGs) and deep learning models have improved the systematic identification of repurposing candidates, most current approaches function as ‘black boxes.’ These comprehensive, state-of-the-art predictors [12] provide predictions without offering clear, human-understandable explanations of their underlying reasoning ([13]). This lack of explainability poses a significant barrier to clinical adoption and validation of computational drug repurposing predictions ([14]). From a biological perspective, the identification of specific pathways connecting drug candidates to disease targets is crucial for several reasons. Biological pathways represent the fundamental mechanisms through which drugs exert their therapeutic effects. Through explainable methods, accurate prediction of pathways provides mechanistic validation that a predicted drug-disease association operates through biologically plausible routes, distinguishing meaningful therapeutic opportunities from spurious correlations ([15]). Moreover, pathway-level explanations enable researchers to assess safety implications by identifying potential off-target effects or conflicting regulatory mechanisms that might contraindicate use in specific patient populations ([16]) ([17]). Additionally, a drug repurposed for a disease can be unrelated to the indication it was initially studied for, as exemplified by Metformin’s evolution from a diabetes treatment to potential anti-cancer properties ([18]). Thus, mechanistic explanations for each prediction are crucial for building clinical trust in AI.

On the other hand, existing predictors that leverage knowledge graphs to predict drug-disease interactions, and give pathway level interpretability, struggle to balance predictive power with plausible explanations. Some of these approaches rely on predefined path schemas [19], limiting their ability to discover novel biological mechanisms. Others use complex learning strategies (such as reinforcement learning) [20] [21] that require extensive tuning, making them less adaptable across different diseases. Rule-based methods provide clear explanations [22] but struggle with generalization, while deep learning based predictors often prioritize accuracy at the cost of transparency [23]. These challenges highlight the need for approaches that can provide biologically meaningful explanations without sacrificing predictive performance.

Apart from the trade-off between performance and explainability, another major challenge that predictors face is data scarcity. Around 92% of 17,080 examined diseases have no approved indications and this scarcity is even more pronounced for rare diseases, where approximately 95% lack FDA-approved treatments and up to 85% have no promising drug candidates ([7], [24]). This limited data constrains predictive methods, leaving them with too few examples to learn meaningful drug-disease relationships. These minimal therapeutic options and limited molecular understanding poses a significant challenge for drug-repurposing predictors ([7]).

These challenges highlight the need for predictors that can effectively learn from sparse data while providing mechanistic insights to validate repurposing predictions, particularly when exploring novel therapeutic applications. To address this gap, we introduce COMIC, a predictor that utilizes **CO**ntrastive **M**asking with **I**nterpretable **C**onnections and aims to balances predictive performance with interpretability. The primary motivation behind incorporating contrastive masking is to identify and preserve the most informative features within sparse data. Through contrastive learning, these networks distinguish meaningful patterns and paths, allowing the predictor to capture both common and rare but significant interaction signatures. This selective approach makes the prediction process interpretable at multiple levels: from identifying critical interaction features through the feature selection network to discovering relevant biological pathways through the path selection network. This enables tracking how each prediction emerges through selected features and pathways while maintaining strong predictive performance. The proposed predictor is evaluated across nine disease categories of the PrimeKG knowledge graph (comprising 17,080 diseases, and 4M+ relationships). COMIC demonstrated a significant performance boost over the current state-of-the-art [7], achieving an average improvement of 4.29% in indication prediction and 14.82% in contraindication prediction, resulting in an overall average gain of 9.55% across both interaction types.

To validate COMIC’s real-world applicability, we evaluated its performance on 30 FDA-approved (2022-2025) repurposed indications that are not present in PrimeKG. The predictor successfully predicted 21 out of 30 (70%) approved indications, with 10 predictions ranking above the 90th percentile. Compared to the state-of-the-art [7], COMIC showed superior discrimination ability. These results demonstrate COMIC’s practical utility while highlighting opportunities for improvement in challenging domains like oncology and neurodegenerative diseases. To facilitate broad adoption, we have deployed our predictor as a web application that provides both predictions and their comprehensive explanations for indications, contraindications, and off-label use https://drp.opendfki.de/.

## 2 Related Work

Recent technological advances have led to the rapid development of numerous knowledge graph-based drug repurposing predictors. These predictors can be categorized into two distinct groups: (1) predictors that solely identify interactions between drugs and diseases and (2) predictors that not only make interaction predictions but also provide explanations. As discussed in the previous section, explanations play a crucial role in drug repurposing; therefore, this section only provides a high-level overview of the second category of predictors. These explainable predictors have evolved into distinct streams, each employs different strategies for model interpretability, ranging from attention-based path weighting to rule-based reasoning systems. While these methods have made significant progress in explaining drug-disease associations, they involve trade-offs between interpretability, computational efficiency, and biological plausibility. **KGML-xDTD** employs a hybrid approach combining knowledge graph embeddings with reinforcement learning to identify drug-disease associations ([19]).

**MIN-ERVA** and **PoLo** employ similar reinforcement learning agents to traverse KGs, generating rule-guided paths that mimic human hypothesis generation ([21]) ([20]). While KMGL-xDTD’s attention-weighted metapaths capture biological relationships, it is constrained by predefined metapath schemas and reinforcement learning based predictors require extensive reward engineering and may overfit to specific traversal patterns, especially on large graphs like PrimeKG.

**XG4Repo** employs the **RNNLogic+’s** framework which uses neural logical reasoning in which a RNN generates logic rules for drug repositioning patterns ([25]) ([26]). Hybrid frameworks like **SAFRAN** integrate KG embeddings with rule-based systems, using Noisy-OR functions to aggregate metapath scores ([22]). These rules based system are often constrained by the expressiveness of their generated rules, which may fail to generalize across diverse drug-disease relationships.

A promising approach involves using graph explanation models to interpret Graph Neural Networks (GNNs). These methods analyze GNNs by selectively masking parts of the graph and assessing how the absence of these parts impacts predictions, thereby highlighting the most influential components in the decision-making process. **RD-Explainer** adapts GNNExplainer to generate semantic subgraphs highlighting genes, symptoms, and pathways ([27]) ([28]). **TxGNN** integrates message-passing graph neural networks with metric learning to rank drug-disease pairs across 17,080 diseases ([7]), achieving state-of-the-art performance. However, its interpretability relies on a post-hoc GraphMask-based explainer to identify critical pathways ([29]). COMIC builds upon TxGNN’s extensive disease coverage and GNN architecture, introducing a contrastive masking-based predictor that provides end-to-end interpretability. Unlike TxGNN, COMIC learns to distinguish relevant features and pathways during training, making interpretability an inherent part of its process. This design allows COMIC to jointly optimize for both prediction accuracy and explanation quality, adapting to complex feature interactions while maintaining strong performance across the full disease spectrum.

## 3 Materials and Methods

This section provides a detailed overview of the COMIC predictor’s architecture, describes the dataset used for its evaluation, and outlines the metrics employed to assess both its predictive performance and the interpretability of its explanations.

### 3.1 The Proposed Predictor

As illustrated in Figure 1, COMIC takes a knowledge graph G as input, which is formally represented as a collection of triples, where each triple t is defined as:

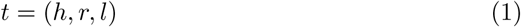

**Fig. 1.**
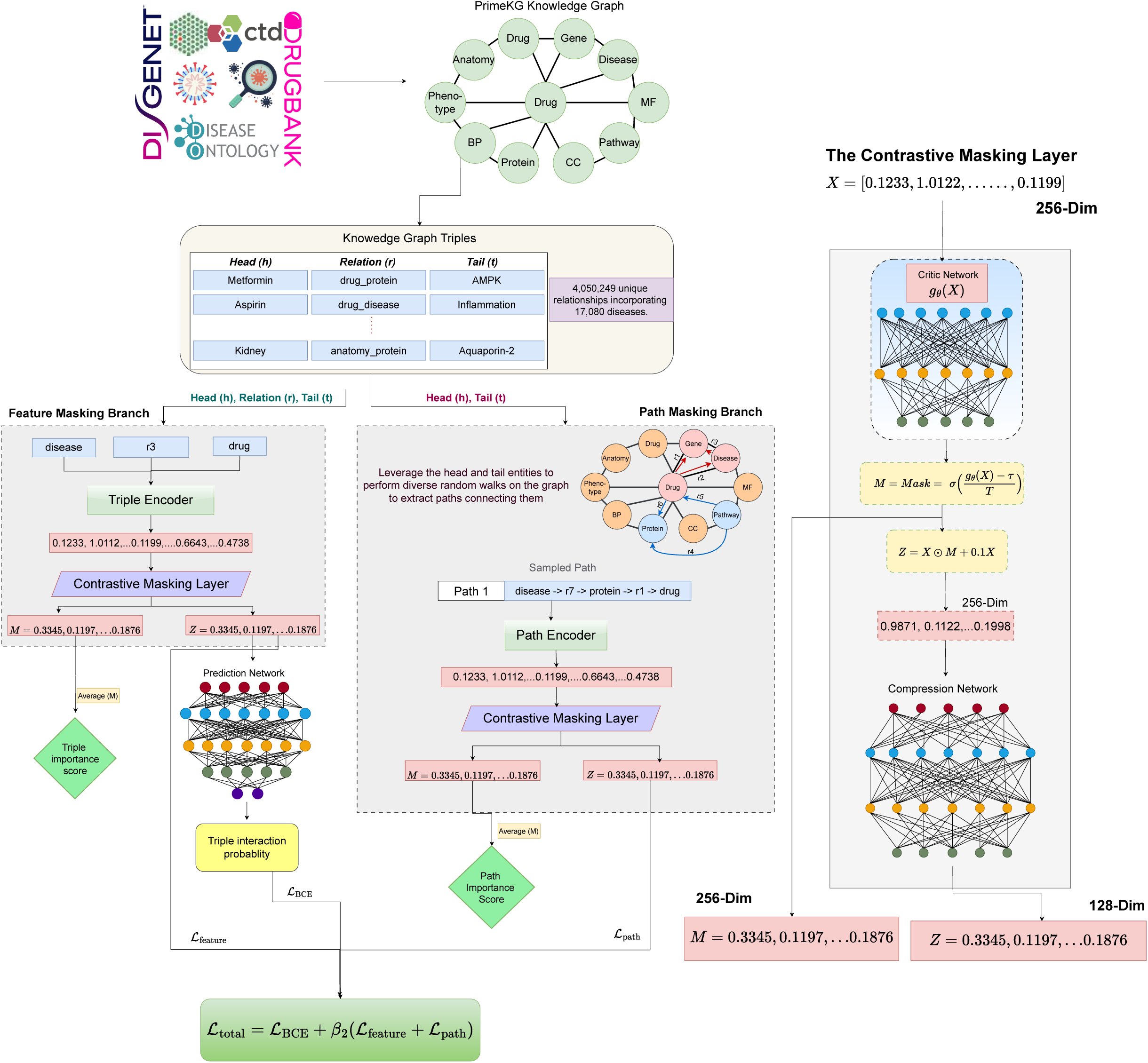
Overview of the COMIC architecture, illustrating the constrastive masking approach

where *h* and *l* denote the head and tail entities respectively, and r represents the relation between them.

The first step is embedding generation. For each entity and relation, dense vector representations are generated:

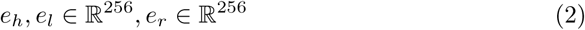

These embeddings are generated through Xavier initialization and are normalized for stability [30]. The normalized entity and relation embeddings then flow into a multi-channel architecture which has two branches: the feature masking branch and the path masking branch.

#### 3.1.1 Feature Masking Branch

This branch first processes each input triple through the Triple Encoding layer and its output is fed into the Feature Masking layer.

The triple encoding layer first concatenates the head, relation, and tail embeddings to preserve their sequential structure. Positional encoding is then applied to enhance this sequence with ordering information. The position-enhanced sequence is passed through a self-attention layer, enabling the triple embedding to maintain the sequence of the triple elements. The attended representation undergoes mean and max pooling, followed by concatenation and compression to produce a 256-dimensional triple representation:

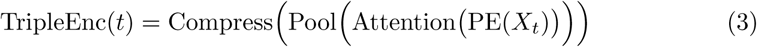

Here, *X_t_*represents the triple input which is comprised of the head, tail, and relation embeddings, and PE denotes positional encoding.

The output of the triple encoding layer is input to a contrastive masking layer that identifies and preserves the most relevant features. The inner working of this layer is described in Section 3.3.

#### 3.1.2 Path Masking Branch

In parallel with triple processing, we samples paths between the head and tail entities (path sampling is detailed in Section 4.3).

Each sampled path is processed by a Path Encoding layer, which embeds each triple in the path using a position-aware neural network. The encoder applies selfattention across all triplets in the path and compresses the attended representation to 256 dimensions. The position-aware encoding ensures the path embedding maintains the sequential nature of the path:

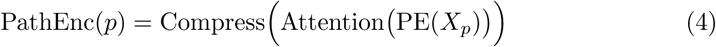

Here, *X_p_*represent the sequence of triplets in the path, and PE denotes positional encoding.

The output of the path encoding layer is input to a contrastive masking layer that identifies and preserves the most relevant biological pathways. The inner working of this layer is described in Section 3.3.

#### 3.1.3 The Contrastive Masking Layer

The Contrastive Masking layer is inspired by the information bottleneck principle of learning compressed, relevant representations. This layer serves as a learnable filter that extract the most relevant features from its input for prediction.

For a given input representation X (either TripleEnc(*X_t_*) or PathEnc(*X_p_*)), the masking layer produces both a filtered representation Z and a selection mask M through:

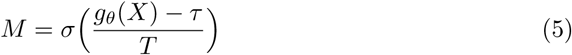

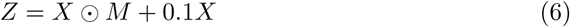

where:

- *g_θ_* is a multi-layer neural network
- *τ* is a learnable threshold
- T is a temperature parameter
- ⊙ represents element-wise multiplication

The selection mask M controls which features are retained, while the residual term 0.1X ensures gradient flow. *g_θ_*is a 3-layer neural network with layer normalization and GELU activation ([31]).

The training objective is to learn a filtered representation Z that preserves only the information relevant for prediction from the original representation X while discarding noise. We accomplish this through contrastive learning using the InfoNCE estimator ([32]). The key idea is to minimize the mutual information between input X and filtered representation Z (compression), while maximizing the mutual information between Z and target label Y. To quantify these information relationships, the InfoNCE estimator estimates mutual information between two representations, X and Z, through contrastive learning:

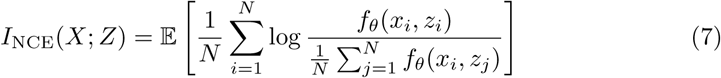

where *f_θ_*is a neural critic function that scores the similarity between the input X and its filtered representation Z, and N is the batch size.

The masking layer processes the encoded representations to produce:

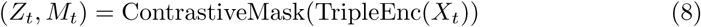

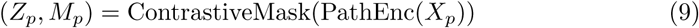

These filtered representations and their corresponding masks enable both prediction and interpretation of the predictor’s decisions. The masks *M_t_* and *M_p_* highlight which features and paths were most relevant for the prediction, while *Z_t_* and *Z_p_* contain the information relevant for prediction, which is used for optimizing the training objective.

The contrastive learning objectives are computed as:

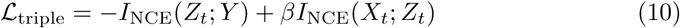

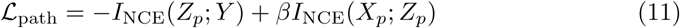

Where Y are the target labels. The negative term maximizes relevant selected information, while the positive term minimizes redundant information.

These losses guide the Contrastive Masking layer to learn optimal feature masks, balancing between retaining relevant information for prediction while discarding noise.

#### 3.1.4 Interaction Prediction and Loss Computation

The filtered triple representations from the feature masking layer (*Z_t_*) are passed through a binary classification neural network. The prediction is computed as:

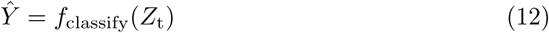

where *f*_classify_ is the classification network (a 3-layer GELU activated neural network).

The Binary Cross Entropy loss between the true labels Y and predicted outputs *Y*^^^ is calculated as:

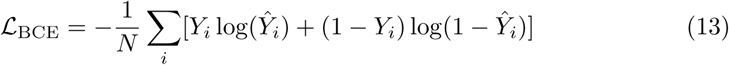

The total loss combines the prediction loss with the contrastive learning objectives:

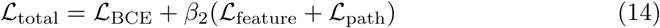

### 3.2 Dataset

Drug repurposing—the process of identifying new therapeutic uses for existing drugs—requires comprehensive understanding of biological interactions across multiple scales, from molecular mechanisms to clinical outcomes. While several biomedical knowledge graphs exist, including SPOKE ([33]), GARD ([34]), BioKG ([35]), and PrimeKG ([10]), we evaluate COMIC on PrimeKG due to its unique characteristics that facilitate in depth understanding of drug-disease relationships. With 22,236 disease terms consolidated into 17,080 clinically meaningful conditions, PrimeKG offers unprecedented disease coverage—1-2 orders of magnitude greater than previous datasets ([10]). This extensive disease representation is crucial for identifying novel drug applications across a wide spectrum of conditions. Furthermore, PrimeKG features 30 distinct relation types specifically designed for drug-related information, including critical relationships for repurposing such as indication, contraindication, and off-label use ([10]). These relations provide complementary signals: indication relationships serve as positive examples of validated therapeutic applications, contraindications highlight important safety boundaries and potential adverse effects, and off-label use cases offer valuable hints about promising therapeutic directions that warrant further investigation. This rich characterization of drug-disease interactions enables COMIC to learn from both successful and unsuccessful therapeutic applications while identifying promising candidates that mirror existing off-label use patterns. The remaining specialized relations enable learning indirect drug-disease interactions, which allows for more contextualized predictions and explanations.

The integration of data from 20 authoritative sources, including DrugBank ([36]), DisGeNET ([37]), CTD ([38]), Gene Ontology ([39]), Human Phenotype Ontology ([40]), and Reactome ([41]), provides PrimeKG with the multi-scale biological context necessary for explainable drug repurposing ([10]). This comprehensive integration manifests in a graph structure comprising 129,375 nodes across 10 biologically relevant types, connected by 4,050,249 unique relationships, capturing the complex web of biological interactions relevant to drug repurposing, from molecular pathways to clinical manifestations.

Additionally, with 90.8% of Orphanet’s 9,348 rare diseases represented, PrimeKG enables drug repurposing for rare diseases—a particularly valuable application given the limited treatment options for these conditions ([10]). These characteristics make PrimeKG an ideal foundation for developing explainable AI systems for drug repurposing, capable of both identifying novel therapeutic applications and providing mechanistic explanations for their predictions.

### 3.3 Evaluation Metrics

To comprehensively evaluate COMIC, we employ two distinct sets of evaluation strategies: one for assessing prediction performance and another for interpretability assessment. For prediction performance, we follow standard practices from the biomedical literature [7][19][28] [42] [43] [44], using Area Under the Precision-Recall Curve (AUPRC) as the primary metric, complemented by accuracy for detailed performance analysis.

The prediction evaluation metrics are summarized in Equation 15:

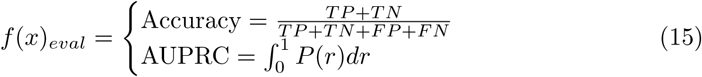

where TP, TN, FP, and FN represent true positives, true negatives, false positives, and false negatives respectively. TPR denotes the true positive rate, FPR the false positive rate, and P(r) the precision at recall value r. These metrics collectively provide a robust assessment of our predictor’s ability to correctly predict drug-disease associations.

Following the evaluation criteria of explainable AI (XAI) literature [45], we assess graph-based explanations using insertion and deletion faithfulness tests, which evaluate the importance of the graph’s edges in the prediction process. Additionally, we assess biological plausibility by comparing the highlighted pathways with established mechanisms of action from medical literature.

We evaluate graph interpretability using the following two metrics: **a. Insertion (faithfulness)**: we start from an EMPTY graph and gradually INSERT top K% edges to the graph where the importance is measured by COMIC’s interpretable connections. We measure the updated AUPRC on the test set using different values of K. A better XAI method has a higher AUPRC at each threshold since it suggests that the newly added edges are informative to the graph predictions (also referred to as fidelity+). **b. Deletion (faithfulness)**: we start from the FULL graph and gradually DELETE top K% edges to the graph where the importance is measured by COMIC’s interpretable connections. We measure the updated AUPRC on the test set using different values of K. A better XAI method has a lower AUPRC at each threshold since it suggests that the deleted edges are necessary for the graph predictions (referred to as fidelity-).

To evaluate the biological plausibility of the pathways highlighted by COMIC, we turn to medical literature and seek pathways that incorporate the indications’ mechanism of action (MOA). Specifically, we conduct a literature review to identify established biological pathways associated with each drug’s therapeutic indications and compare these with our model’s highlighted pathways.

## 4 Experimental Setup

This section covers the proposed predictor’s experimental and implementation details. COMIC is implemented in PyTorch 2.0 for GPU-accelerated computation, with Numpy for handling vectorized operations during preprocessing and inference phases ([46]) ([47]). The web application is implemented in Django with a PostgreSQL backend.

### 4.1 Data Splits

We utilize the data split methodology established by Huang et al., which splits PrimeKG into nine disease areas [7]. These nine splits were constructed to mirror realworld drug repurposing scenarios where new diseases have limited associated data [7]. Each holdout set is carefully curated following three strict criteria: complete exclusion of approved drugs from the training data, minimal disease similarity with the training set, and removal of biological neighbor information to limit molecular data availability [7]. Given all diseases in a given disease area, all indications and contraindications were removed from the dataset used to train machine learning models. Additionally, a fraction (5%) of the connections between biomedical entities to these diseases were removed from PrimeKG [7]. The splits encompass major therapeutic domains including adrenal gland disorders, autoimmune diseases, neurodegenerative conditions, metabolic disorders, and cardiovascular diseases, with additional splits covering anemia, diabetes, cancer, and mental health. This systematic partitioning from Huang et al. [7] ensures COMIC is tested under realistic drug repurposing conditions, where the predictor must generalize to diseases with limited prior knowledge.

### 4.2 Negative Sampling Strategies

Negative sampling is a pivotal technique in Graph based Learning, essential for training predictors to effectively distinguish between valid and invalid relationships. The quality of negative samples significantly influences the performance of predictors, as they provide the necessary contrast to positive samples during training. However, generating high-quality negative samples poses challenges, particularly in ensuring that these samples are informative and contribute to robust learning. [48]. Various negative sampling strategies have been proposed in the literature to address these challenges. Random negative sampling is the most straightforward approach, where entities in triples are randomly replaced to create negative samples. While efficient, this method can produce trivial negatives that may not effectively challenge the predictor [48]. Model-guided and knowledge-constrained strategies leverage pre-trained models or external knowledge to generate semantically meaningful negatives, albeit with increased computational complexity [48]. Moreover, probabilistic negative sampling adjusts the sampling probability based on entity frequency, focusing on harder negatives.

In our work, we implement two distinct negative sampling strategies:

1. **Degree-Based Hard Negative Sampling**: This approach generates negative samples by replacing either the head or tail entity in a triple, with the replacement probability proportional to the node degrees. Higher-degree nodes have a higher likelihood of being selected as replacements, resulting in more challenging negative samples that can enhance training by focusing on harder-to-predict cases.
2. **Random Sampling with Fixed Destination Nodes**: In this strategy, negative samples are created by maintaining the destination nodes while randomly selecting source nodes from the set of valid entities. To ensure diversity and robustness in learning, we adopt an epoch-wise regeneration strategy, where new random negative samples with fixed destinations are generated at the beginning of each training epoch.

### 4.3 Path Sampling Approach

In large-scale knowledge graphs, comprehensive path finding between entities is often impractical due to computational constraints [49]. To address this, various path sampling strategies have been developed. One common approach is the use of meta-paths, which are predefined sequences of relation types that capture specific semantic relationships within heterogeneous graphs. This method leverages domain knowledge to guide the sampling process, focusing on meaningful and interpretable connections [50]. Alternatively, random walk-based sampling offers a more flexible approach by exploring the graph without any constraints. This allows for the discovery of novel and potentially unexpected connections, as the predictor learns path patterns directly from the data.

In our study, we implement a random walk sampling strategy to explore the knowledge graph:

1. **Random Walk Sampling**: For each drug-disease pair, we perform random walks to sample up to 5 distinct paths, each limited to a maximum length of 5 relations. Then, 5 false paths are created by randomly corrupting the last two nodes from each path.
2. **Diversity Threshold**: To ensure the quality of extracted evidence, we implement a diversity threshold that prevents the collection of redundant paths. Specifically, a new path is accepted only if its overlap with previously sampled paths falls below 50%, compelling the sampler to identify varying patterns between each drug-disease pair.

### 4.4 Training Procedure

We train the COMIC predictor using the Adam optimizer [51] with a learning rate of 0.0005. The predictor is trained for a maximum of 30 epochs with a batch size of 1024. For the contrastive masking layer, we set the temperature parameter to start at 1.0 and anneal it linearly throughout training to a minimum of 0.1, while *β* is fixed at 0.2. We employ early stopping with a patience of 2 epochs on the validation loss to prevent overfitting and select the best performing weights. All hyperparameters are tuned with the validation set, and we use gradient clipping with a maximum norm of 1.0 to ensure stable training.

## 5 Results

This section illustrates the COMIC predictor’s performance under four distinct experimental settings. (1) A comprehensive comparison of COMIC against eight baseline methods and TxGNN using the established disease area splits. (2) Validation of COMIC’s real-world applicability by testing its predictions on recent FDA approvals that occurred after PrimeKG’s construction. (3) Assessment of COMIC’s interpretability through insertion and deletion experiments that demonstrate the predictor’s ability to identify biologically meaningful relationships. (4) Examination of the biological credibility of COMIC’s explanatory pathways through case studies of recently approved drug-disease pairs.

### 5.1 Predictive Performance

COMIC’s predictive performance is compared against eight baseline methods and TxGNN using the disease area splits and results reported by [7] (described in Section 4.2.1). The baseline methods span network statistical techniques (Kullback–Leibler divergence, Jensen–Shannon divergence), graph-theoretical approaches (network proximity, diffusion state distance), modern GNN architectures (RGCN, HGT, HAN), and language models (BioBERT) [52] [53] [54] [17] [55] [56] [57] [58]. To ensure robust comparison, we first conducted an analysis of negative sampling strategies. While Huang et al.[7] employed both random and degree-based sampling in their work, they did not specify which approach was used for their final reported results. Given this ambiguity, we evaluated both approaches through validation on recent FDA-approved drugs. We found that degree-based hard negative sampling outperformed random generation by an average margin of 10.2% in ranking correct indications higher. This superior performance better reflects real-world drug repurposing scenarios by forcing the predictor to correctly distinguish between therapeutically similar but distinct drug-disease pairs. More importantly, by choosing degree-based hard negative sampling for our comparative analysis, we ensure our reported performance metrics represent conservative estimates relative to potentially simpler random sampling approaches. All our results reported in this section use degree-based hard negative sampling and are averaged across five random seeds (1-5), providing a robust and conservative comparison baseline.

COMIC demonstrated consistently high performance across all nine disease areas for both indication and contraindication prediction tasks (Table 1 and 2). For indications, COMIC achieved AUPRC scores ranging from 0.88 to 0.94, surpassing the state-of-the-art (Table 1). Notably, while the state-of-the-art [7] showed strong performance in specific areas (AUPRC 0.95 for adrenal gland, anemia, and neurodegenerative diseases), COMIC maintained more consistent performance across all disease categories, with an average improvement of 4.29% over TxGNN. This consistency is particularly evident in challenging areas like cardiovascular diseases, where COMIC achieved an AUPRC of 0.88 compared to TxGNN’s 0.64, representing a 37.8% improvement. For contraindications, COMIC outperformed existing methods across all disease areas, with AUPRC scores ranging from 0.88 to 0.96 (Table 2). The predictor achieved particularly strong results in diabetes (AUPRC: 0.95) and metabolic disorders (AUPRC: 0.95), areas where accurate contraindication prediction is crucial for patient safety. The performance gap was again most pronounced in cardiovascular diseases, where COMIC maintained robust performance (AUPRC: 0.88) while other methods showed significant degradation. Again, COMIC maintained more consistent performance across all disease categories, with an average improvement of 14.82% over TxGNN. COMIC overall achieved an average gain of 9.55% across both interaction types. These results establish COMIC’s predictive power in a test setting.

**Table 1.** Performance in predicting drug indications. Reported is AUPRC.

| Category | KL | JS | Proximity | DSD | RGCN | HAN | HGT | BioBERT | TxGNN | COMIC |
| --- | --- | --- | --- | --- | --- | --- | --- | --- | --- | --- |
| random | 0.50 | 0.49 | 0.53 | 0.49 | 0.84 | 0.87 | 0.72 | 0.81 | 0.91 | 0.9375 |
| cell proliferation | 0.51 | 0.50 | 0.47 | 0.51 | 0.47 | 0.65 | 0.61 | 0.83 | 0.92 | 0.9031 |
| mental health | 0.50 | 0.50 | 0.44 | 0.50 | 0.55 | 0.61 | 0.56 | 0.64 | 0.89 | 0.9046 |
| cardiovascular | 0.49 | 0.50 | 0.50 | 0.50 | 0.41 | 0.63 | 0.60 | 0.66 | 0.64 | 0.8817 |
| anemia | 0.47 | 0.53 | 0.49 | 0.51 | 0.42 | 0.67 | 0.53 | 0.64 | 0.96 | 0.9302 |
| adrenal gland | 0.48 | 0.50 | 0.48 | 0.50 | 0.38 | 0.59 | 0.54 | 0.64 | 0.99 | 0.9375 |
| autoimmune | 0.51 | 0.49 | 0.56 | 0.52 | 0.51 | 0.53 | 0.49 | 0.66 | 0.87 | 0.9284 |
| metabolic disorder | 0.51 | 0.54 | 0.41 | 0.51 | 0.67 | 0.56 | 0.54 | 0.68 | 0.76 | 0.9362 |
| diabetes | 0.50 | 0.52 | 0.47 | 0.52 | 0.40 | 0.67 | 0.62 | 0.69 | 0.87 | 0.9114 |
| neurodegenerative | 0.50 | 0.51 | 0.49 | 0.51 | 0.84 | 0.71 | 0.60 | 0.57 | 0.95 | 0.9182 |

**Table 2.** Performance in predicting drug contraindications. Reported is AUPRC.

| Category | KL | JS | Proximity | DSD | RGCN | HAN | HGT | BioBERT | TxGNN | COMIC |
| --- | --- | --- | --- | --- | --- | --- | --- | --- | --- | --- |
| random | 0.49 | 0.50 | 0.56 | 0.49 | 0.77 | 0.84 | 0.63 | 0.58 | 0.82 | 0.9646 |
| cell proliferation | 0.49 | 0.49 | 0.50 | 0.50 | 0.54 | 0.51 | 0.47 | 0.78 | 0.87 | 0.9233 |
| mental health | 0.50 | 0.50 | 0.50 | 0.50 | 0.67 | 0.44 | 0.56 | 0.54 | 0.79 | 0.9267 |
| cardiovascular | 0.49 | 0.49 | 0.51 | 0.50 | 0.63 | 0.47 | 0.51 | 0.56 | 0.72 | 0.8836 |
| anemia | 0.50 | 0.50 | 0.50 | 0.50 | 0.65 | 0.54 | 0.51 | 0.53 | 0.74 | 0.9497 |
| adrenal gland | 0.49 | 0.50 | 0.48 | 0.49 | 0.75 | 0.46 | 0.55 | 0.47 | 0.88 | 0.9502 |
| autoimmune | 0.50 | 0.49 | 0.51 | 0.50 | 0.68 | 0.40 | 0.61 | 0.40 | 0.86 | 0.9486 |
| metabolic disorder | 0.49 | 0.51 | 0.47 | 0.51 | 0.58 | 0.56 | 0.51 | 0.53 | 0.78 | 0.9525 |
| diabetes | 0.50 | 0.51 | 0.48 | 0.51 | 0.57 | 0.56 | 0.52 | 0.51 | 0.69 | 0.9564 |
| neurodegenerative | 0.51 | 0.51 | 0.56 | 0.51 | 0.71 | 0.42 | 0.55 | 0.60 | 0.78 | 0.9560 |

### 5.2 Evaluation of COMIC on recent FDA-approvals

The next step is to validate COMIC’s real-world applicability. For this, we test the predictor on 30 repurposed drug-disease pairs that received FDA approval between 2022-2025. These approvals occurred after the knowledge graph’s construction, ensuring that no relationships between these drug-disease pairs existed in the training data. Highlighted in Table 3, this evaluation provides a stringent test of the predictor’s ability to identify drug repurposing candidates.

**Table 3.**
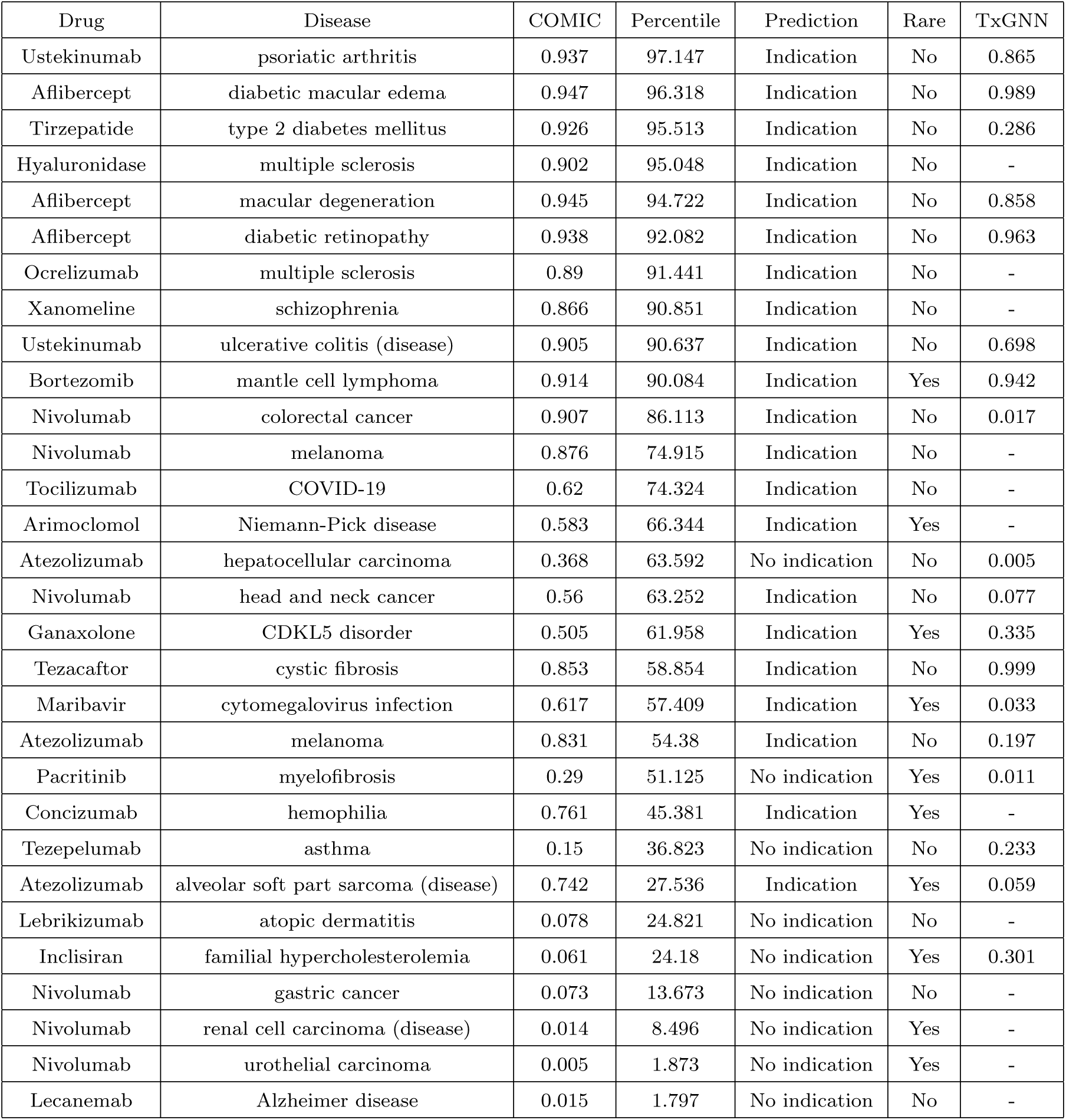
COMIC’s performance on recent FDA-Approved Repurposed Indications. The percentile indicates how high the drug was ranked relative to all possible indications for the disease.

Despite achieving exceptional AUPRC scores on the test set, COMIC’s performance on completely novel FDA approvals revealed important real-world challenges. The predictor successfully predicted 21 out of 30 (70%) approved indications, with 10 of these predictions ranking above the 90th percentile (Table 3). This notable performance gap between testing and prospective evaluation underscores the inherent difficulty of predicting therapeutic applications, particularly when mechanistic understanding is still evolving for most diseases.

COMIC still demonstrated strong predictive ability for several therapeutic areas. For autoimmune conditions, Ustekinumab for Psoriatic Arthritis ranked in the 97th percentile, while for metabolic disorders, Tirzepatide for Type 2 Diabetes ranked in the 95th percentile. Aflibercept showed consistently high rankings across multiple indications: diabetic macular edema (96th percentile), macular degeneration (94th percentile), and diabetic retinopathy (92nd percentile).

To further investigate the gap in performance, we compared COMIC’s performance against TxGNN using the TxGNNExplorer (txgnn.org) [7], which provided TxGNN scores for 18 out of the 30 drug-disease pairs as shown in Table 3. The comparison reveals COMIC’s superior discrimination ability (scoring higher for 12 out of 18 pairs), suggesting our predictor captures complex molecular interactions better in the context of these pairs.

Where COMIC struggled most was with certain cancers and novel therapeutic mechanisms. For instance, Nivolumab received varied predictions across different cancer types, ranging from high confidence for Colorectal Cancer (86th percentile) to low confidence for Urothelial Carcinoma (1.8th percentile) and Renal Cell Carcinoma (8.5th percentile). Similarly, recent Alzheimer’s disease therapy Lecanemab received a low ranking (1.8th percentile), highlighting challenges in neurodegenerative diseases where their mechanisms are still being understood [59]. These results reveal both COMIC’s strengths and limitations in real-world drug repurposing scenarios. While the 70% prediction rate demonstrates substantial practical utility, the performance differential between the test set and recent approvals highlights opportunities for further refinement.

### 5.3 Evaluation of COMIC’s Interpretability

To assess COMIC’s interpretability, we conduct insertion and deletion evaluation described in Section 3.3. In the insertion experiment, as shown in Figure 3, the AUPRC steadily increased—from 0.67 at 10% insertion up to 0.99 when all edges were inserted—demonstrating that the edges identified as influential indeed contribute substantially to COMIC’s predictive performance. Conversely, the deletion test started with the full graph and gradually removed edges ranked by their importance. As shown in Figure 2, the resulting deletion curve reveals a marked decrease in AUPRC at each step (from approximately 0.99 with no removal to 0.62 at 90% deletion), highlighting that even modest removals of highly ranked edges significantly impair COMIC’s outputs.

**Fig. 2.**
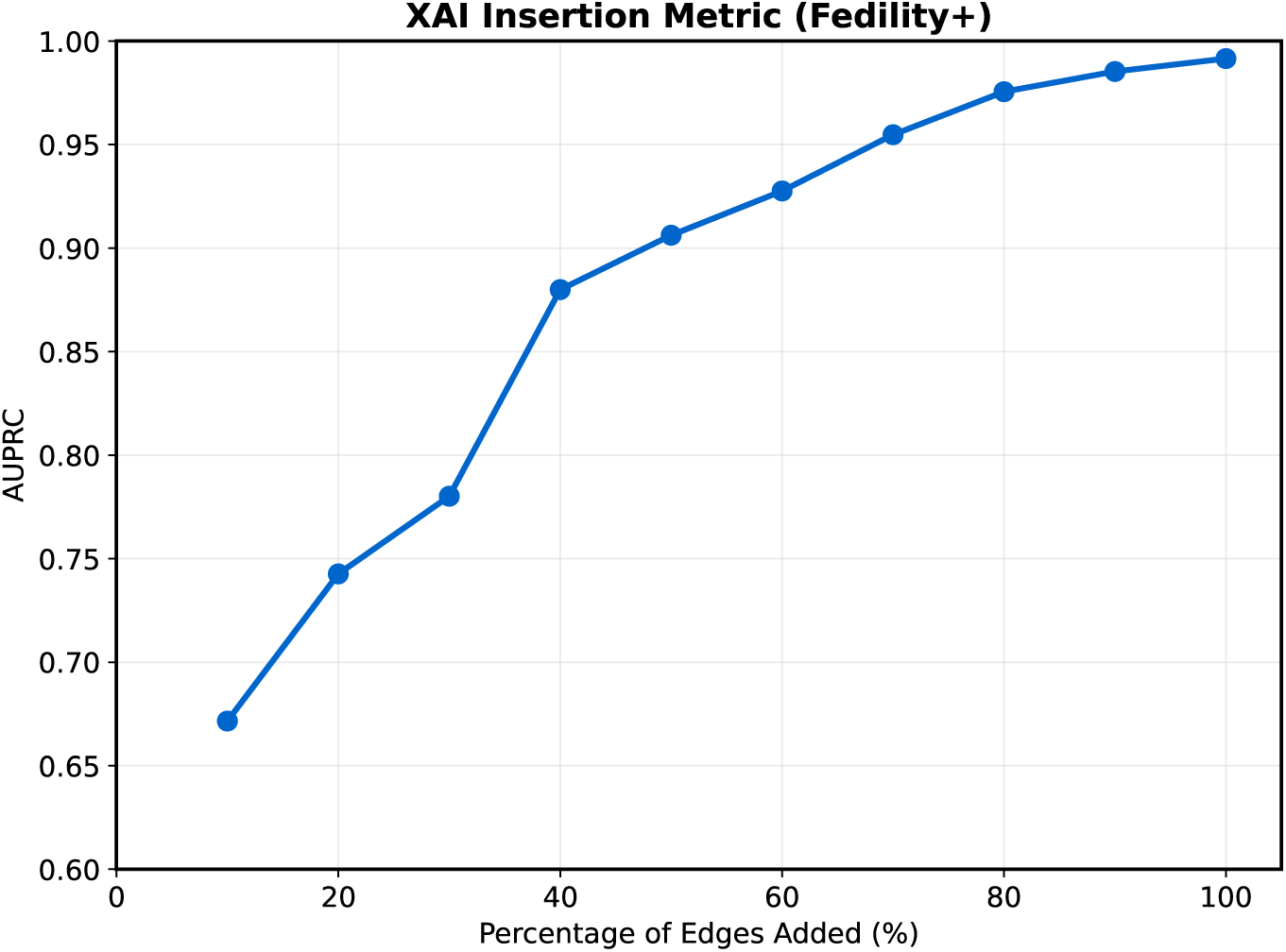
Insertion Performance Evaluation

**Fig. 3.**
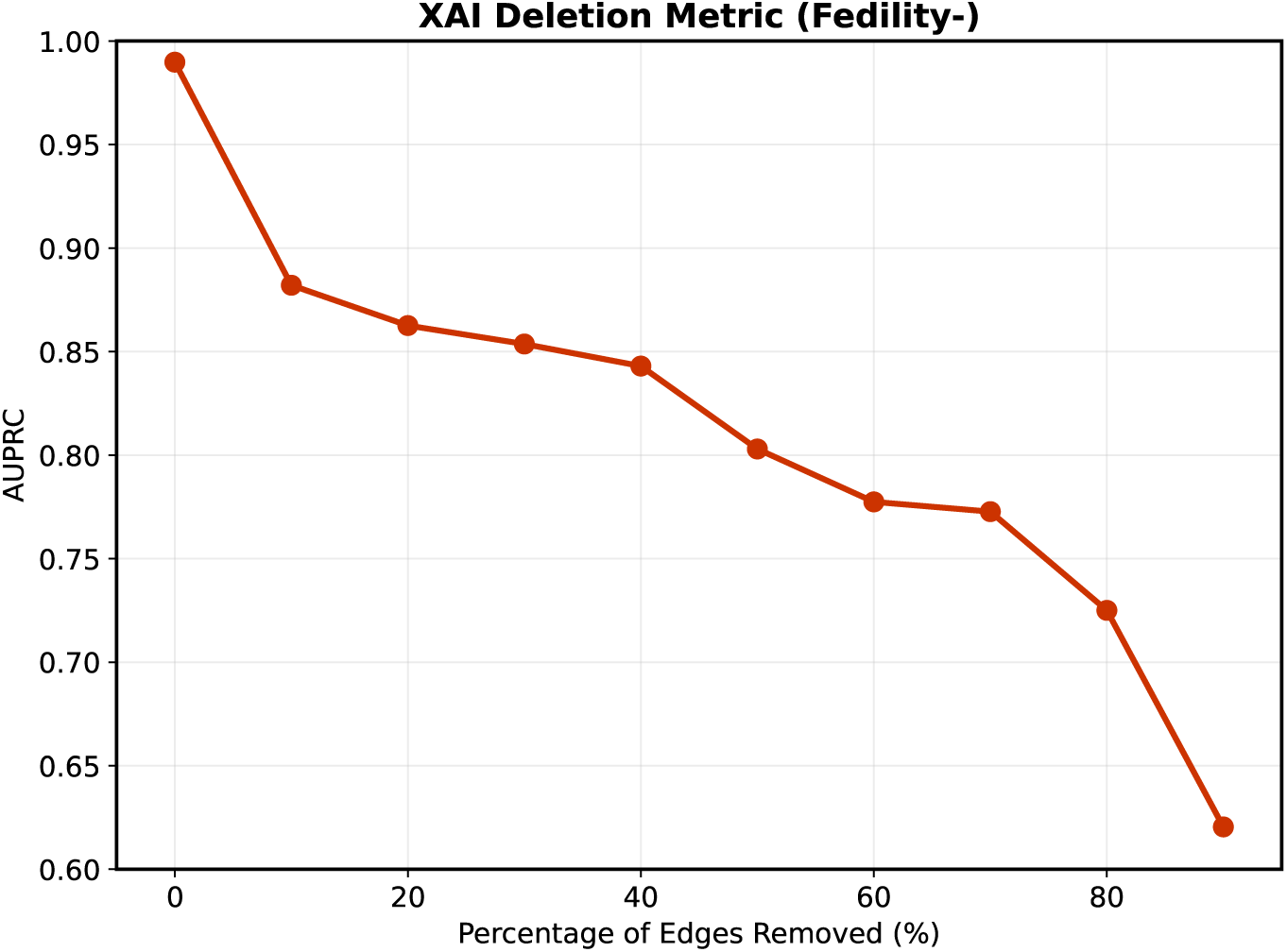
Deletion Performance Evaluation

The initial steep decrease in deletion and the initial steep increase in insertion conform to the ideal in these evaluation metrics, confirming that COMIC’s interpretable connections are faithful to the predictor’s underlying decision-making process. Together, these trends indicate that the most highly-ranked edges are truly critical for accurate predictions as indicated by the rapid initial degradation during deletion and high initial gains during insertion. This demonstrates that COMIC’s interpretability is transparent and robust in identifying the edges most crucial to the predictor’s decision process.

### 5.4 Mechanistic Pathways Validation

We now examine the biological credibility of its explanatory pathways. We consider the indication pairs in Table 3 for stringent testing since they were approved after PrimeKG’s construction.

In the case of Ocrelizumab for Multiple Sclerosis, COMIC identified several pathways that directly link its target, MS4A1 (CD20), to key immune mediators ([60]). Notably, several valid pathways traverse through molecules such as HLA-DRA, CD40, NF-KB, and IL7R—elements that are well documented in the pathogenesis of multiple sclerosis and align closely with the drug’s known MOA of B cell depletion and subsequent modulation of T cell responses ([61]) ([62]) ([63]). These mechanistic explanations are further supported by literature that underscores the importance of MHC class II interactions and costimulatory signals in mediating autoimmunity ([64]). Similarly, for Atezolizumab in Melanoma, COMIC revealed pathways that directly connect CD274 (PD-L1) to known melanoma drivers ([65]). In particular, pathways such as the MTOR–AKT1 cascade reflect established oncogenic processes in melanoma ([66]). Additionally, a link through SLC39A11 to PMEL (gp100) directly ties to a melanocyte differentiation antigen and a key target in melanoma immunotherapy ([67]). These paths provide mechanistically plausible explanations that resonate with current understanding of melanoma biology and the rationale behind immune checkpoint inhibition.

In contrast, for Bortezomib and Mantle Cell Lymphoma, the predictor predominantly returned paths based on drug effects—for instance, adverse effects such as dyssynergia, hyperuricemia, connect to drugs, which then eventually tie to other phenotypes like fatigue and, indirectly, to mantle cell lymphoma. Such paths do not capture the primary mechanism of action of Bortezomib, but instead appear to be driven by indirect associations present in the dataset. In cases like this where the underlying molecular mechanisms are not well represented in the knowledge graph, COMIC did not find any meaningful mechanistic pathways. For instance, for Alzheimer’s disease and lecanemab, for asthma and tezepelumab, or for hemophilia and concizumab, the predictor’s explanations were either absent or based on indirect disease associations rather than direct MOA-related pathways.

These examples underscore that while COMIC is capable of retrieving highly plausible mechanistic paths when the underlying biology is present in the dataset, its explanations for indications lacking comprehensive mechanistic data tend to rely on indirect associations. This reliance on indirect associations when insufficient evidence is not in the dataset for certain indications emphasizes the need for further enrichment of the underlying knowledge graph and continued refinement of our interpretability methods.

### 5.5 The COMIC Web Application

After extensive evaluation of COMIC’s capabilities, we have deployed the predictor as a web application (https://drp.opendfki.de/) that provides predictions and their explanations for drug indications and contraindications. Users can select/search diseases, configure pathway parameters, and search for specific drugs. Results are presented in a sortable table with confidence scores for each drug-disease pair. The application also visualizes the pathway explanations connecting drugs to diseases through interactive network diagrams, helping researchers understand the potential mechanisms of action. This web applications aims to accelerate drug repurposing research by making COMIC’s predictions accessible to the broader scientific community, serving as a valuable resource for drug development researchers, clinicians exploring alternative therapies, and computational biologists investigating disease mechanisms.

## 6 Conclusion

The core contribution of this work lies in the development of a novel predictor, COMIC, that enables the systematic identification of potential new indications for existing drugs, with a particular focus on diseases that are characterized by a lack of effective therapeutic options. This approach offers a significant advancement in addressing the unmet medical needs associated with these conditions. COMIC achieves a 4.29% average improvement in indication prediction over the state-of-the-art in a test setting. The superior performance of the COMIC predictor, compared to existing approaches, is driven by its innovative feature and path masking branches. The feature masking branch successfully identifies critical drug-disease interaction signatures, while the path masking branch is capable of highlighting mechanistic pathways, together providing clinicians with transparent reasoning for COMIC’s predictions. COMIC’s real-world applicability is validated through its correct identification of 70% of recently approved therapeutic indications, demonstrating considerable practical utility while maintaining end-to-end interpretability. But the evaluation on recent FDA approvals also revealed important limitations, particularly for diseases with evolving molecular understanding such as certain cancers and neurodegenerative conditions. These challenges highlight the critical need for richer knowledge graphs with more comprehensive mechanistic data, especially for rare diseases where treatment options remain limited. Future research should focus on expanding the biological pathway representations within these graphs and refining interpretable mechanisms to better capture complex therapeutic relationships. By addressing these data gaps and continuing to enhance interpretability, we can further bridge the divide between computational predictions and clinical adoption of AI-driven drug repurposing.

## Supplementary information

Accuracy results on train, valid, and test sets of each of the nine disease splits have been attached as supplementary material.

## Declarations

- **Funding:** There is no funding source.
- **Conflict of interest/Competing interests:** There is no author’s conflict of interest.
- **Ethics approval and consent to participate:** Research is conducted responsibly, with respect for participants’ autonomy, dignity, and well-being
- **Consent for publication:** Authors provide consent for their work to be shared in journal.
- **Data availability:** The dataset supporting the conclusions of this article is available in the Harvard Dataverse repository. And the data splits can be accessed from the TxGNN GitHub Repository.
- **Materials availability:** All material is available in the paper.
- **Code availability:** The predictor is publicly available at the following URL: https://drp.opendfki.de/
- **Authors contribution**:
  – **Naafey Aamer:** Conceptualization, Methodology, Validation, Investigation, Formal analysis, Visualization, Data curation, Manuscript writing
  – **Muhammad Nabeel Asim:** Conceptualization, Methodology, Validation, Manuscript writing, review and editing manuscript, Supervision.
  – **Andreas Dengel:** Supervision, Reviewed the article and performed final editing

